# Employing NaChBac for cryo-EM analysis of toxin action on voltage-gated Na^+^ channels in nanodisc

**DOI:** 10.1101/2020.04.16.045674

**Authors:** Shuai Gao, William C Valinsky, Nguyen Cam On, Patrick R Houlihan, Qian Qu, Lei Liu, Xiaojing Pan, David E. Clapham, Nieng Yan

**Author notes:** To whom correspondence should be addressed: N. Yan.

## Abstract

NaChBac, the first bacterial voltage-gated Na^+^ (Na_v_) channel to be characterized, has been the prokaryotic prototype for studying the structure-function relationship of Na_v_ channels. Discovered nearly two decades ago, the structure of NaChBac has not been determined. Here we present the cryo-EM analysis of NaChBac in both detergent micelles and nanodiscs. Under both conditions, the conformation of NaChBac is nearly identical to that of the potentially inactivated Na_v_Ab. Determining the structure of NaChBac in nanodiscs enabled us to examine gating modifier toxins (GMTs) of Na_v_ channels in lipid bilayers. To study GMTs in mammalian Na_v_s, we generated a chimera in which the extracellular fragment of the S3 and S4 segments in the second voltage-sensing domain from Na_v_1.7 replaces the corresponding sequence in NaChBac. Cryo-EM structures of the nanodisc-embedded chimera alone and in complex with HuwenToxin IV (HWTX-IV) were determined to 3.5 Å and 3.2 Å resolutions, respectively. Compared to the structure of HWTX-IV-bound human Na_v_1.7, which was obtained at an overall resolution of 3.2 Å, the local resolution of the toxin has been improved from ~ 6 Å to ~ 4 Å. This resolution enabled visualization of toxin docking. NaChBac can thus serve as a convenient surrogate for structural studies of the interactions between GMTs and Na_v_ channels in a membrane environment.

---

Voltage-gated sodium (Na_v_) channels initiate and propagate action potentials by selectively permeating Na^+^ across cell membranes in response to changes in the transmembrane electric field (the membrane potential) in excitable cells (1–3). There are nine subtypes of Na_v_ channels in humans, Na_v_1.1-1.9, whose central role in electrical signaling is reflected by the great diversity of disorders that are associated with aberrant functions of Na_v_ channels. To date, more than 1000 mutations have been identified in human Na_v_ channels that are related to epilepsy, arrhythmia, pain syndromes, autism spectrum disorder, and other diseases (4–6). Elucidation of the structures and functional mechanisms of Na_v_ channels will shed light on the understanding of fundamental biology and facilitate potential clinical applications.

The eukaryotic Na_v_ channels are comprised of a pore-forming α subunit and auxiliary subunits (7). The α subunit is made of one single polypeptide chain that folds to four repeated domains (I-IV) of six transmembrane (S1-S6) segments. The S5 and S6 segments from each domain constitute the pore region of the channel, which are flanked by four voltage sensing domains (VSDs) formed by S1-S4. The VSD, a relatively independent structural entity, provides the molecular basis for voltage sensing in voltage-dependent channels (Na_v_, K_v_, and Ca_v_ channels) and enzymes (3, 8–10).

Owing to the technological advancement of electron cryomicroscopy (cryo-EM), we were able to resolve the atomic structures of Ca_v_ and Na_v_ channels from different species and in complex with distinct auxiliary subunits, animal toxins, and FDA-approved drugs (11–20). Among these, human Na_v_1.7 is of particular interest to the pharmaceutical industry for the development of novel pain killers; mutations in Na_v_1.7 are associated with various pain disorders, such as indifference to pain or extreme pain syndrome (21–28). The structures of human Na_v_1.7 bound to the tarantula gating modifier toxins (GMTs) in combination with pore blockers (Protoxin II (ProTx-II) with tetrodotoxin (TTX); Huwentoxin IV (HWTX-IV) with saxitoxin (STX)), in the presence of β1 and β2 subunits were determined at overall resolutions of 3.2 Å (17). The GMTs, which associate with the peripheral regions of VSDs, were only poorly resolved as ‘blobs’ that prevented even rigid-body docking of the toxin structures.

The poor resolution of the toxins may be due to the loss of interaction with the membrane in detergent micelles; intact membranes were reported to facilitate the action of these toxins (29–32). Understanding the binding and modulatory mechanism of GMTs may require structures of the channel-toxin complex in a lipid environment, such as nanodiscs. However, the low yield of Na_v_1.7 recombinant protein has impeded our effort to reconstitute the purified protein into nanodiscs. We therefore sought to establish a surrogate system to facilitate understanding the modulation of Na_v_1.7 by GMTs.

NaChBac from *Bacillus halodurans* was the first prokaryotic Na_v_ channel to be identified (33). More orthologs of this family were subsequently characterized (34). In contrast to eukaryotic counterparts, bacterial Na_v_ channels are homo-tetramers of 6-TM subunits. Bacterial Na_v_ channels, exemplified by Na_v_Ab and Na_v_Rh, have been exploited as important model systems for structure-function relationship studies (35–38). Despite having been discovered two decades ago, the structure of the prototypical NaChBac remains unknown.

The characterized GMT binding sites are not conserved between prokaryotic and eukaryotic Na_v_ channels. Bacterial Na_v_ channel-based chimeras have been employed for structural analysis of the mechanisms of GMTs and small molecule perturbation. In these chimeras, the extracellular halves of VSDs were replaced by the corresponding sequences from the eukaryotic counterparts. These studies have afforded important insight into the function of GMTs at the VSD and have facilitated the structural elucidation of Na_v_ channels in distinct conformations (39, 40).

The most commonly used scaffold protein is Na_v_Ab; however, overexpression and purification of Na_v_Ab in insect cells is costly and slow. Alternatively, Na_v_Rh cannot be recorded in any tested heterologous system (36). Therefore, we focused on NaChBac. By overcoming technical challenges with recombinant NaChBac suitable for cryo-EM analysis, we were able to solve the structures of NaChBac in detergent micelles and in nanodiscs. A NaChBac chimera that is able to bind to HWTX-IV was generated; its structures alone and in complex with HWTX-IV in nanodiscs were determined to 3.5 Å and 3.2 Å resolutions, respectively.

## Results

### Cryo-EM analysis of NaChBac in detergents and in nanodisc

Since its discovery in 2001, each of our laboratories made various attempts to obtain structural information for NaChBac. Although the channel can be overexpressed in *E. coli* BL21(DE3) in mg/l LB culture and can be recorded in multiple heterologous systems, it tends to aggregate during purification, preventing crystallization trials or cryo-EM analysis.

After extensive trials for optimization of constructs, expression and purification conditions, the Yan lab determined the key to obtaining a mono-dispersed peak of NaChBac on size exclusion chromatography (SEC). At pH 10.0, NaChBac is compatible with several detergents, with glyco-diosgenin (GDN) being the best of those tested (Figure S1). We purified NaChBac in the presence of 0.02% (w/v) GDN and solved its structure to 3.7 Å out of 91,616 selected particles using cryo-EM (Figures S2, S3A, C, D, Table S1).

We then sought to reconstitute NaChBac into nanodiscs. Preliminary trials revealed instability of the nanodisc-enclosed NaChBac at pH 10.0. We modified the protocol and lowered the pH to 8.0 when first incubating the protein with lipids. Excellent samples were obtained after these procedures (Figure 1A), and the structure of NaChBac in nanodiscs was obtained at 3.1 Å resolution out of 281,039 selected particles (Figures S2-S4, Table S1). For simplicity, we will refer to the structures obtained in the presence of GDN and nanodiscs as NaChBac-G and NaChBac-N, respectively.

**Figure 1.**
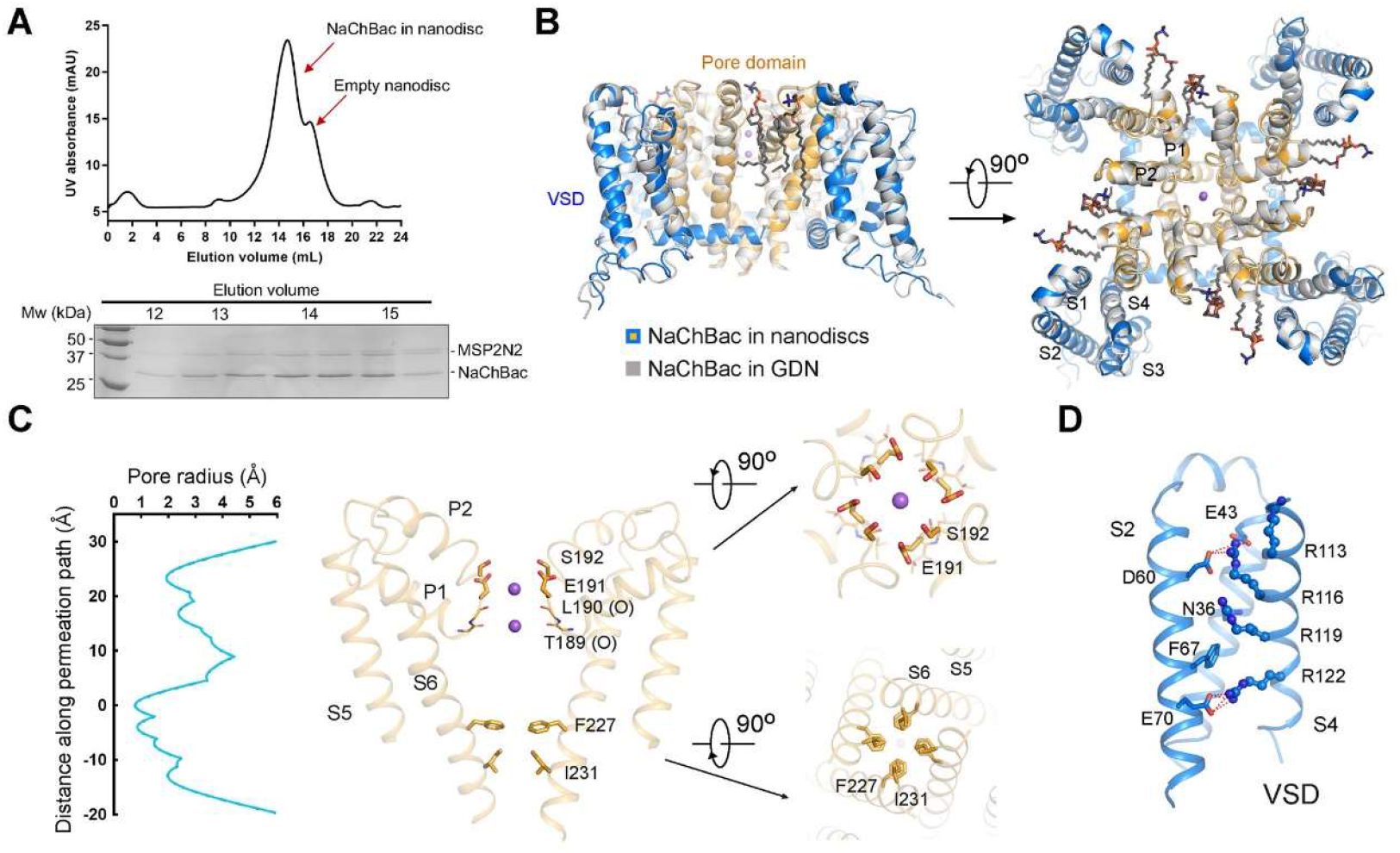
Structural determination of NaChBac in GDN micelles and in nanodiscs. **(A)** Final purification step for NaChBac in MSP2N2-surrounded nanodiscs. Shown here is a representative size-exclusion chromatogram (SEC) and Coomassie blue-stained SDS-PAGE. **(B)** Structures of NaChBac in GDN and nanodiscs are nearly identical. The structure of NaChBac-N (in a nanodisc) is colored orange for the pore domain and blue for VSDs, and that of NaChBac-G (gray in GDN) is colored gray. Two perpendicular views of the superimposed structures are shown. The bound ions and lipids are shown as purple spheres and black sticks, respectively. **(C)** Permeation path of NaChBac-N. The pore radii, calculated by HOLE (47), are shown on the left and the selectivity filter (SF) and the intracellular gate are shown on the right. **(D)** Structure of the VSD. Three gating charge residues are above the occluding Phe67 and one reside below. All structural figures were prepared in PyMol (48).

### Structure of NaChBac in a potentially inactivated state

Despite different pH conditions and distinct surrounding reagents, the structures of NaChBac remained nearly identical in GDN and in nanodiscs, with a root-mean-square deviation of 0.54 Å over 759 Cα atoms for the tetrameric channel (Figure 1B). For both structures, application of a four-fold symmetry led to improved resolution, indicating identical conformations of the four protomers. Because of the higher resolution, we will focus on NaChBac-N for structural illustrations. In both reconstructions, two lipids are found to bind to each protomer (Figure 1B).

Two spherical densities can be resolved in the EM map and half maps in the selectivity filter (SF) of NaChBac-N in both C1 and C4 symmetry, supporting the assignment of two ions (Figure S5). Although we cannot exclude the possibility of contaminating ions, we will still refer to the bound ions as Na^+^, which was present at 150 mM throughout purification. As observed in the structures of other homotetrameric Na_v_ channels, an outer Na^+^ coordination site is enclosed by the side groups of Glu191 and Ser192 and an inner site is constituted by the eight carbonyl oxygen groups from Thr189 and Leu190 (Figure 1C, Figure S4B).

Calculation of the pore radii with the program “HOLE” (41) reveals that the intracellular gate, formed by Ile231 on S6, is closed (Figure 1C). Note that Phe227, which is one helical turn above Ile231, has ‘up’ and ‘down’ conformations that were resolved in the 3D EM reconstruction (Figure S4C). In the “down” state, it is part of the closed gate, whereas in the “up” conformation it no longer contributes to the constriction (Figure 1C). These dual conformations of Phe at the intracellular gate is similar to those in the structures of human Na_v_1.7 and Na_v_1.5 (17, 20).

In each VSD, S4 exists as a 3_10_ helix. Three gating charge (GC) residues on S4, Arg113/116/119, are above, and one, Arg122, is below the occluding Phe67 on S2 (Figure 1D). The overall structure, which is similar to the first reported structure of Na_v_Ab (PDB code: 3RVY), may represent a potentially inactivated conformation.

### Structures of NaChBac-Na_v_1.7VSD_II_ chimera alone and bound to HWTX-IV

Inspired by the successful structural resolution of NaChBac in nanodiscs, we then attempted to generate chimeras to confer GMT binding activity to NaChBac. The linker between S3 and S4 on the second VSD (VSD_II_) of Na_v_1.7 has been characterized to be the primary binding site for HWTX-IV (42). For one construct, we substituted the NaChBac sequence (residues 98-110) comprising the extracellular halves of S3 and S4 and their linker with the corresponding segment from the human Na_v_1.7-VSD_II_ (residues 817-832); we refer to this as the chimera (Figure 2A).

**Figure 2.**
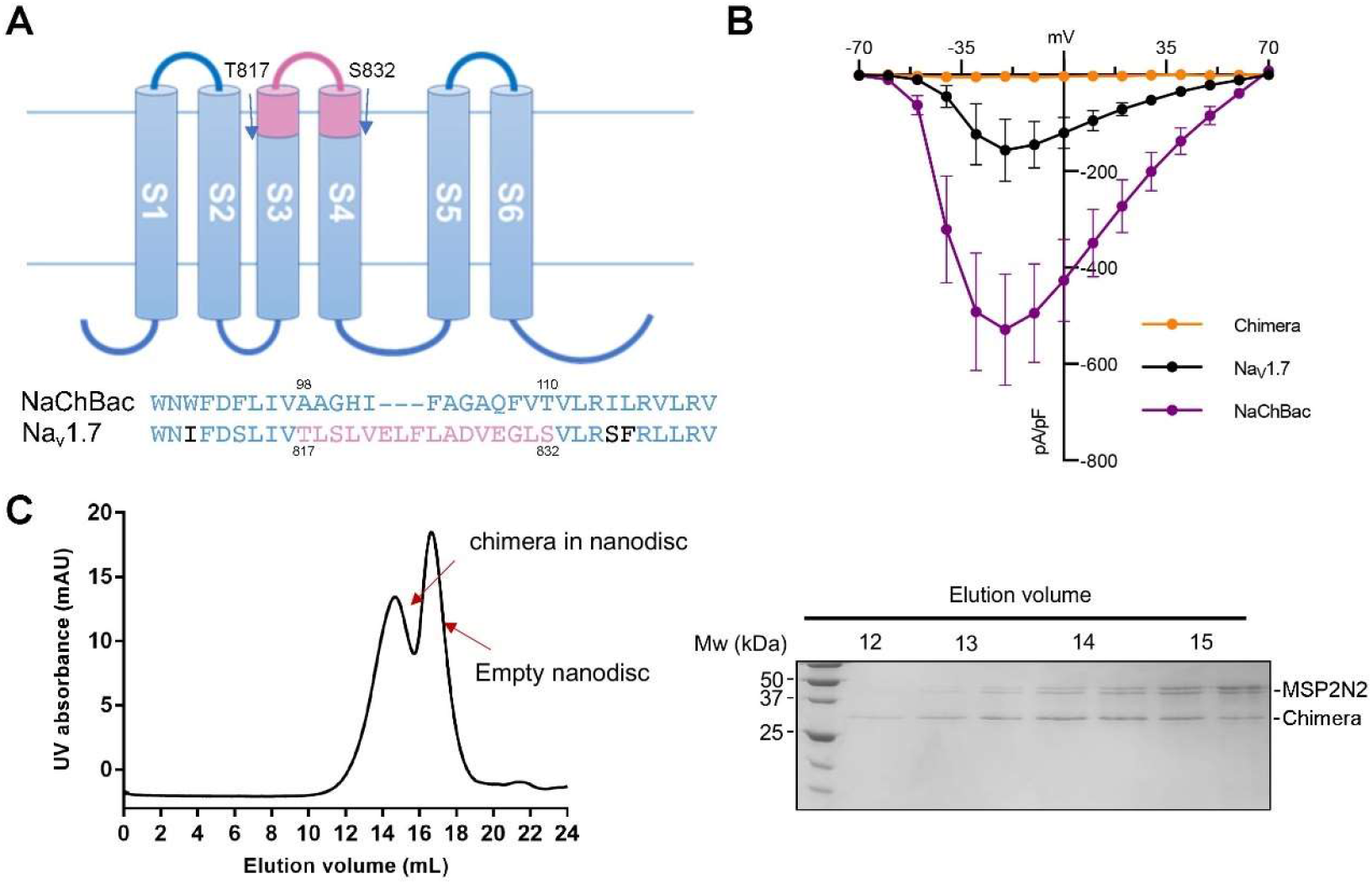
Engineering of a NaChBac-Na_v_1.7VSDII chimera. **(A)** A diagram for the chimera design. Residues 98-110 of NaChBac are replaced by the corresponding sequence, colored pink, from human Na_v_1.7. **(B)** Current-voltage relationships of the chimera (orange; n = 7), Na_v_1.7 (black; n = 6), and NaChBac (purple; n = 7) elicited from a step protocol from −70 mV to +70 mV (in 10 mV increments). Macroscopic currents were not reliably detected for the chimera. **(C)** Final SEC purification of the chimera in the POPC nanodisc. Coomassie blue-stained SDS-PAGE for the indicated fractions are shown on the right.

Although both NaChBac and Na_v_1.7 can be recorded when expressed in HEK-293 cells, and the chimera maintained decent protein yields from *E. coli* overexpression and biochemical behavior in solution, currents could not be recorded from the chimera (Figure 2B, C, Figure S6). This is not a rare problem for eukaryotic and prokaryotic Na_v_ chimeras (39, 40, 43). Nevertheless, we proceeded with cryo-EM analysis of the chimera alone and in the presence of HWTX-IV.

After some trials, the chimera alone was reconstituted into the brain polar lipid nanodiscs and its complex with HWTX-IV was reconstituted in POPC nanodiscs. Their 3D EM reconstructions were obtained at resolutions of 3.5 Å and 3.2 Å, respectively (Figure 3A, B, Figure S2, S3D, Table S1). The overall structure of the chimera is nearly identical to NaChBac except for the region containing the grafted segments. In addition, four additional lipids are clearly resolved in both chimera reconstructions, for a total of 12 lipids (Figure 3C, Figure S7A).

**Figure 3.**
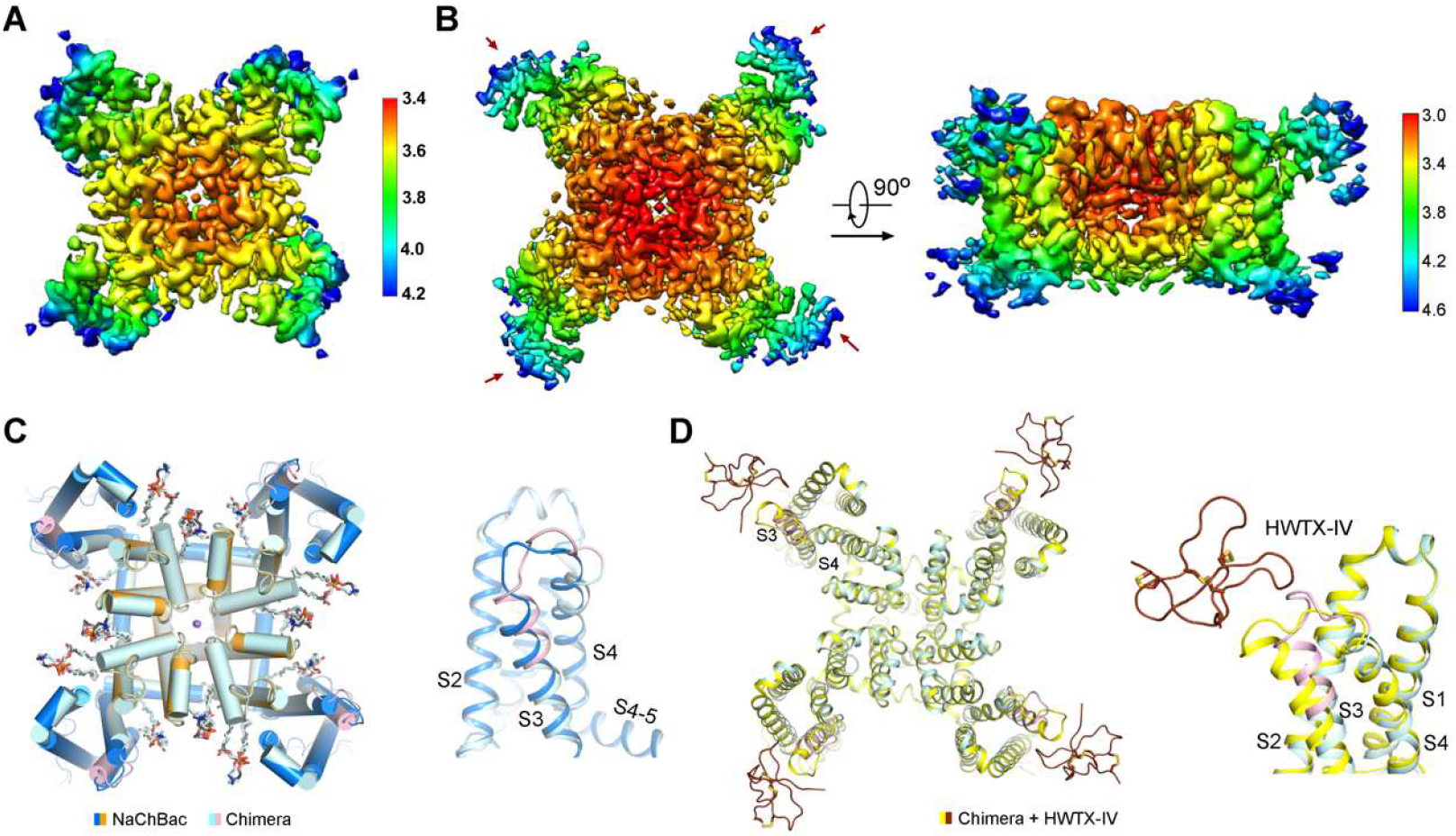
Cryo-EM structures of the chimera alone and in complex with HWTX-IV. **(A, B)** Resolution maps for the apo (A) and HWTX-IV-bound chimera (B). The resolution maps were calculated with Relion 3.0 (49) and prepared in Chimera (50). **(C)** Structural shift of the chimera upon HWTX-IV binding. Left: extracellular view of the superimposed NaChBac-N and chimera. Right: conformational deviation of the grafted region in the VSD. **(D)** HWTX-IV attaches to the periphery of VSD. The S3-S4 loop moves toward the cavity of the VSD upon HWTX-IV binding.

The densities corresponding to HWTX-IV are observed on the periphery of each VSD, with a local resolution of 4.2-4.6 Å (Figure 3B). The NMR structure of HWTX-IV (PDB ID: 1MB6) (29) could be docked as a rigid body into these densities, although the side chains were still indiscernible (Figure S7B,C). In the presence of the toxin, the local structure of the Na_v_1.7 fragment undergoes a slight outward shift (Figure 3D). Because of the structural similarity between the chimera and NaChBac-N, we will focus the structural analysis on the interface between the chimera and HWTX-IV.

### HWTX-IV is half-inserted in the lipid bilayer

Although the local resolution of the toxin is insufficient for side chain analysis, its backbone could be traced. Half of the toxin appears to be submerged in the densities corresponding to the nanodisc (Figure 4A). When the two reconstructions are overlaid (Figure 4B), the toxin bound to the chimera is positioned lower (towards the nanodisc) than in the GDN-surrounded human Na_v_1.7 structure. In addition, the S3-S4 loop of VSD_II_, which was not resolved in human Na_v_1.7 (17), is now visible in the chimera (Figure S7A). These differences support the importance of the lipid bilayer in positioning HWTX-IV and its binding segment.

**Figure 4.**
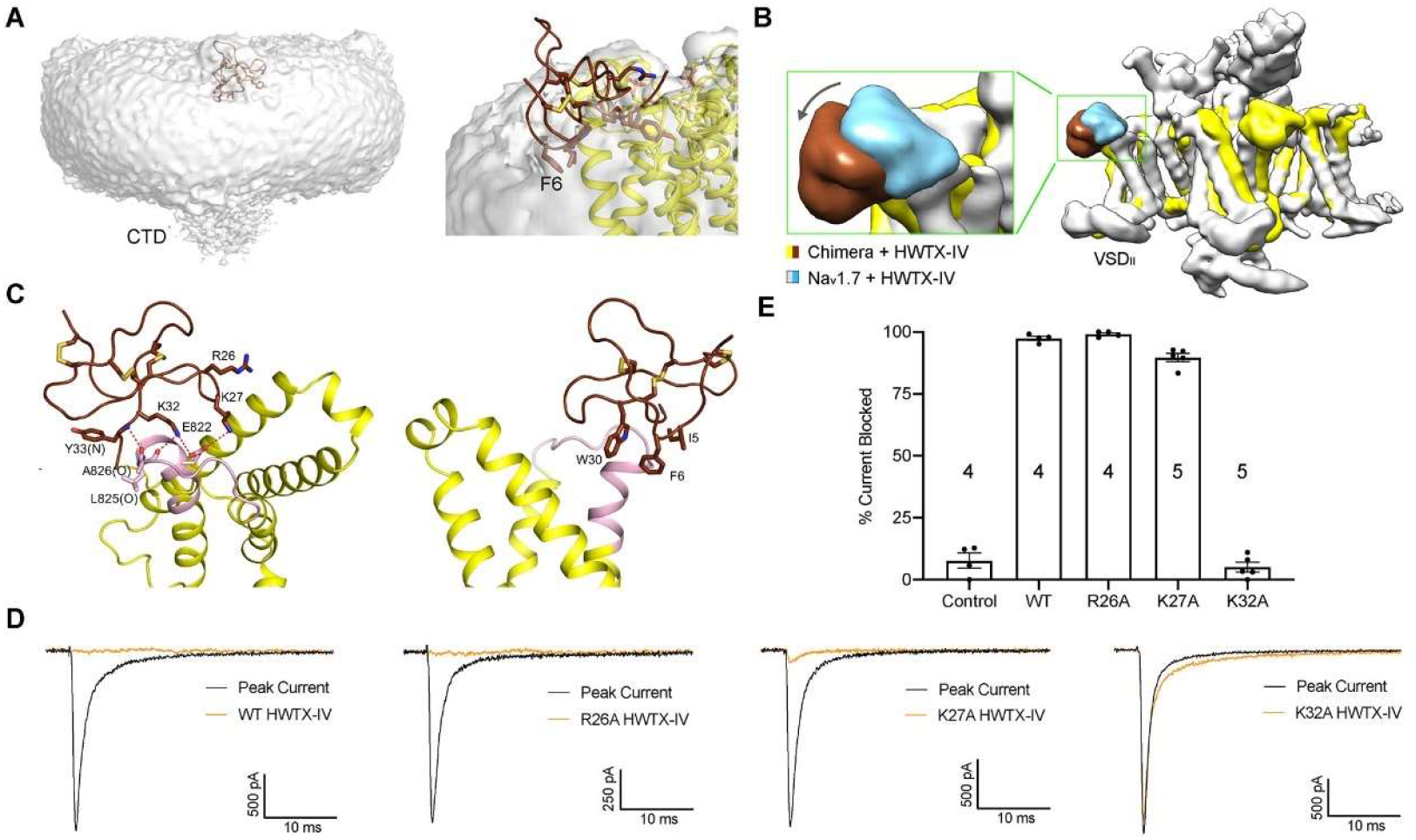
HWTX-IV inserts into the nanodisc. **(A)** HWTX-IV is half-submerged in the nanodisc. The map was low pass-filtered to show the nanodisc contour. Densities that likely belong to the extended S6 segments (labeled as CTD) at low resolution. **(B)** HWTX-IV resides ‘lower’ in the nanodisc-embedded chimera than that in the detergent-surrounded human Na_v_1.7. The 3D EM maps of HWTX-IV-bound Na_v_1.7 (EMD-9781) and the chimera, both low-pass filtered to ~ 8 Å, are overlaid relative to the pore domain in the chimera. **(C)** Potential interactions between HWTX-IV and the chimera. Note that the local resolution does not support reliable side chain assignment of HWTX-IV. Please refer to Figure S6 for the densities of the toxin and the adjacent channel segment. Shown here are the likely interactions based on the chemical properties of the residues. **(D)** Lys32 of HWTX-IV is critical for Na_v_1.7 suppression. Shown here are representative recordings of Na_v_1.7 current block by the indicated HWTX-IV variants, each applied at 200 nM, elicited from a single step protocol from −100 mV (holding) to −20 mV (maximal activation; 50 ms). (E) Percentage current block (mean +/− SEM) by HWTX-IV variants. Control (no toxin; n = 4), WT (n = 4), R26A (n = 4), K27A (n = 5), and K32A (n = 5).

Based on structural docking, a cluster of three hydrophobic residues, Ile5, Phe6, and Trp30 of HWTX-IV are completely immersed in the lipid bilayer. These side chain assignments are supported by two observations. First, among the 35 residues in HWTX-IV, six Cys residues form three pairs of central disulfide bonds to stabilize the toxin structure. Second, the surface of the toxin is amphiphilic, and the polar side of the toxin is unlikely to be embedded in a hydrophobic environment. This residue assignment placed several positively charged residues, including Arg26, Lys27, and Lys32, in the vicinity of the grafted Na_v_1.7 segment for potential polar interactions (Figure 4C).

Supported by the coordination observed in the toxin bound structure, Glu822 and Glu829 in Na_v_1.7 have been reported to be critical for HWTX-IV sensitivity. Crosslinking experiments suggested an important interaction between Na_v_1.7-Glu822 and Lys27 of HWTX-IV (44–46). Based on these structural implications, we substituted the basic residues Arg26, Lys27, and Lys32 with alanine and examined their action on human Na_v_1.7 through electrophysiology (Figure 4D). While R26A has a negligible effect on toxin activity, K27A exhibited slightly reduced inhibition. In contrast, HWTX-IV(K32A) no longer inhibited Na_v_1.7 activation (Figure 4D, E). These characterizations support the placement of basic residues at the HWTX-IV interface with the grafted Na_v_1.7 segment.

Based on chemical principles and these experiments, we fine-tuned the structural model. Lys32 of HWTX-IV interacts with Glu822, and, along with the backbone amide of Tyr33, forms hydrogen bonds with the backbone groups of Leu825 and Ala826, respectively. This also leaves HWTX-IV’s Lys27 close to Glu822 in the Na_v_1.7 chimera, while Arg26 is beyond the range of effective electrostatic interaction (Figure 4C).

## Discussion

Here we report the long-sought structures of NaChBac, the prototypical bacterial Na_v_ channel that has been extensively biophysically characterized. The convenience of overexpressing the recombinant protein in *E.coli* and reconstituting it into nanodiscs made NaChBac a suitable target for high-resolution cryo-EM analysis. We showed that NaChBac can be used as a surrogate to study the specific interactions between modulators and human Na_v_ channels through chimera engineering. The shift of the binding position of the toxin in nanodiscs from that observed in detergent micelles highlights the importance of a lipid environment for structural investigation of the function of GMTs (Figure 4B).

Notwithstanding this progress, we note that the VSDs of the chimera still remain in the same “up” conformation as in the apo-chimera or in NaChBac, although HWTX-IV was expected to suppress activation of Na_v_ channels (42). The lack of functional expression of chimeric channels also represents a drawback for the surrogate system in the establishment of these structure-function relationships. Further optimization of NaChBac chimera is required to make it a more useful surrogate system.

## Materials and Methods

### Expression and purification of NaChBac and Nav1.7-NaChBac chimera

The cDNA for full-length NaChBac was cloned into pET15b with an amino terminal His_6_ tag. For the Na_v_1.7-NaChBac chimera, residues 98-110 of NaChBac were replaced by residues 817-832 from Na_v_1.7 through standard two-step PCR. Over-expression in *E. coli* BL21(DE3) cells was induced with 0.2 mM IPTG (final concentration) at 22°C when OD600 reached 1.0-1.2. Cells were harvested after 16-h incubation at 22°C, and cell pellets were resuspended in the buffer containing 25 mM Tris pH 8.0, and 150 mM NaCl. Cells were disrupted by sonication (1.5 min/liter), and insoluble fractions were removed by centrifugation at 27,000 g for 10 min. The supernatant was applied to ultra-centrifugation at 250,000 g for 1 h. The membrane-containing pellets were resuspended in the extraction buffer containing 25 mM glycine, pH 10.0, 150 mM NaCl, and 2% (w/v) DDM, incubated at 4 °C for 2 h, and subsequently ultra-centrifuged at 250,000 g for 30 min. The supernatant was applied to Ni-NTA resin, washed with 10 column volumes of wash buffer containing 25 mM glycine, pH 10.0, 150 mM NaCl, 25 mM imidazole, and 0.02% GDN. Target proteins were eluted with 4 column volumes of elution buffer containing 25 mM glycine, pH 10.0, 150 mM NaCl, 250 mM imidazole, and 0.02% GDN. After concentration, proteins were further purified with size exclusion chromatography (SEC; Superdex 200 Increase 10/300 GL, GE Healthcare) in running buffer containing 25 mM glycine, pH 10.0, 150 mM NaCl, and 0.02% GDN.

### Nanodisc reconstitution

Lipid (POPC or BPL, Avanti) in chloroform was dried under a nitrogen stream, and resuspended in reconstitution buffer containing 25 mM glycine, pH 10.0, 150 mM NaCl, and 0.7% DDM. Approximately 250 μg purified protein was mixed with 1 mg MSP2N2 and 750 μg lipid (BPL for NaChBac, POPC for chimera), resulting in a protein-MSP-lipid molar ratio of 1: 12: 520. Approximately 100 μg TEV protease was added to the mixture to remove the His tag from MSP2N2. The mixture was incubated at 4 °C for 5 h with gentle rotation. Bio-beads (0.3 g) were then added to remove detergents from the system and facilitate nanodisc formation. After incubation at 4 °C overnight, Bio-beads were removed through filtration, and protein-containing nanodiscs were collected for cryo-EM analysis after final purification by Ni-NTA resin and SEC in running buffer containing 25 mM Tris, pH 8.0, and 150 mM NaCl (Superose 6 Increase 10/300 GL, GE Healthcare).

### Chemical synthesis of HWTX-IV and its mutations

HWTX-IV and its mutations were synthesized on 500 mg Rink Amide-AM resin (Tianjin Nankai HECHENG) (0.5 mmol/g resin) using the microwave peptide synthesizer (CEM Corporation) to generate the C-terminal peptide amide. The resin was swelled in dimethylformamide (DMF, J&K Scientific) for 10 min. The 9-fluorenylmethyloxycarbonyl (Fmoc) protecting groups were removed by treatment with piperidine (10 % v/v) and 0.1 M ethyl cyanoglyoxylate-2-oxime (Oxyma, Adamas-beta) in DMF at 90 °C for 1.5 min. The coupling reaction was carried out using a DMF solution of Fmoc-amino acid (4 equiv, GL Biochem), Oxyma (4 equiv) and N, N’-diisopropylcarbodiimide (DIC, 8 equiv, Adamas-beta) at 90 °C. Each coupling step required 3 min and then the resin was washed x3 by DMF. The peptide chain elongation proceeded until N-terminal glutamic acid coupling. Finally, a cleavage cocktail, which contains trifluoroacetic acid (TFA, J&K Scientific), H_2_O, thioanisole (J&K Scientific), 1,2-ethanedithiol (EDT, J&K Scientific) at a volume ratio of 87.5: 5: 5: 2.5, was used to remove the protecting groups of side chains and cleave the peptide from resin via a 3 h incubation. In the next step, the cleavage solution was collected and concentrated by pure N_2_ superfusion. After precipitation in cold diethyl ether and centrifugation, the crude peptides were obtained. The linear peptide was analyzed and purified by RP-HPLC.

The lyophilized linear peptide (4.1 mg, 1 μmol) was dissolved in 100 ml refolding buffer containing 100 μmol reduced glutathione (GSH, Sigma), 10 μmol oxidized glutathione (GSSG, J&K Scientific) and 10 mM Tris-HCl pH 8.0 at 25 °C overnight. The folded product was monitored and purified by RP-HPLC.

### Cryo-EM data acquisition

Aliquots of 3.5 μl concentrated samples were loaded onto glow-discharged holey carbon grids (Quantifoil Cu R1.2/1.3, 300 mesh). Grids were blotted for 5 s and plunge-frozen in liquid ethane cooled by liquid nitrogen using a Vitrobot MarK IV (Thermo Fisher) at 8 °C with 100 percent humidity. Grids were transferred to a Titan Krios electron microscope (Thermo Fisher) operating at 300 kV and equipped with a Gatan Gif Quantum energy filter (slit width 20 eV) and spherical aberration (Cs) image corrector. Micrographs were recorded using a K2 Summit counting camera (Gatan Company) in super-resolution mode with a nominal magnification of 105,000x, resulting in a calibrated pixel size of 0.557 Å. Each stack of 32 frames was exposed for 5.6 s, with an exposure time of 0.175 s per frame. The total dose for each stack was ~ 53 e^−^/Å^2^. SerialEM was used for fully automated data collection. All 32 frames in each stack were aligned, summed, and dose weighted (51) using MotionCorr2 (52) and 2-fold binned to a pixel size of 1.114 Å/pixel. The defocus values were set from −1.9 to −2.1 μm and were estimated by Gctf (53).

### Image processing

Totals of 2068/1729/1590/ 2423 cryo-EM micrographs were collected, and 949,848/923,618/798,929/1,490,375 particles were auto-picked by RELION-3.0 (54–56) for NaChBac in GDN, nanodiscs, chimera, and chimera-HWTX-IV in nanodiscs, respectively. Particle picking was performed by RELION-3.0 (57). All subsequent 2D and 3D classifications and refinements were performed using RELION-3.0. Reference-free 2D classification using RELION-3.0 was performed to remove ice spots, contaminants, and aggregates, yielding 718,086/569,435/552,252/713,753 particles, respectively. The particles were processed with a global search K=1 or global multi-reference research K=5 procedure using RELION-3.0 to determine the initial orientation alignment parameters using bin2 particles. A published PDB structure of human Na_v_1.7-NavAb chimera was used as the initial reference. After 40 iterations, the datasets from the last 6 iterations (or last iteration) were subject to local search 3D-classifications using 4 classes. Particles from good classes were then combined and re-extracted with a box size of 240 and binned pixel size of 1.114 Å for further refinement and 3D-classification, resulting in 450,478/302,656/87,482/391,361 particles, which were subjected to auto-refinement. 3D reconstructions were obtained at resolutions of 3.9 Å/3.1 Å/3.9 Å/3.7 Å. Skip-alignment 3D classification using bin1 particles yielded data sets containing 91,616/281,039/31,147/49,149 particles respectively, and resulted in respective reconstructions at 3.7 Å/3.1 Å/3.5 Å/3.2 Å by using a core mask (58).

### Model building and structure refinement

The initial model of NaChBac, which was built in SWISS-MODEL (59) based on the structure of Na_v_1.7-VSD4-Na_v_Ab (PDB: 5EK0), was manually docked into the 3.7 Å EM map in Chimera (60) and manually adjusted in COOT (61), followed by refinement against the corresponding maps by the phenix.real_space_refine program in PHENIX (62) with secondary structure and geometry restraints.

A definitive binding pose for the HWTX-IV could be determined; although the density for many sidechains did not allow unambiguous assignment. The NMR structure (PDB ID: 1MB6) was manually docked into the EM map in Chimera and adjusted in COOT.

### Electrophysiological characterizations

Plasmids carrying cDNAs were co-transfected with EGFP (9:1 ratio) into HEK293 cells for 24 h. Cells were selected for patch-clamp based on green fluorescence. Cells were bathed in a solution containing (mM): 140 Na-Gluconate, 4 KCl, 2 CaCl_2_, 1 MgCl_2_, 10 glucose, and 10 HEPES; pH 7.4. Thick-walled borosilicate patch pipettes of 2 MΩ to 5 MΩ contained (mM): 130 CsCH_3_SO_3_, 8 NaCl, 2 MgCl_2_, 2 MgATP, 10 EGTA, 20 HEPES; pH 7.2. Solutions were 290 +/− 5 mOsm.

Whole-cell currents were recorded at room temperature using an Axopatch 200B patch-clamp amplifier controlled via a Digidata 1440A (Molecular Devices) and pClamp 10.3 software. Data were digitized at 25 kHz and low-pass-filtered at 5 kHz. Cells were held at −100 mV and stepped from −70 mV to +70 mV in +10 mV increments (1000 ms step duration for NaChBac/chimera; 50 ms step duration for Na_v_1.7). After each step, cells were briefly stepped to −120 mV (50 ms) before returning to the holding potential (−100 mV). The inter-sweep duration was 6.2 s for Na_v_1.7 and 15 s for NaChBac. Before recordings, currents were corrected for pipette (fast) capacitance, whole-cell capacitance, and series resistance compensated to 80%. Leak currents and capacitive transients were minimized using online P/4 leak subtraction.

For HWTX-IV recordings, cells were held at −100 mV and stepped to −20 mV (maximal activation) for 50 ms (Na_V_1.7) or 1000 ms (NaChBac). After each step, cells were stepped to −120 mV as described above. The inter-sweep duration, capacitance correction, series resistance compensation, and P/4 leak subtraction were identical to the I-V protocol described above. To study HWTX-IV, the toxin was diluted to the working concentration in extracellular solution and bath-perfused at a rate of 1-2 ml for up to 15 min. Where block was observed, maximum block typically occurred between 10 and 15 min of perfusion. Prior to HWTX-IV perfusion, cells were recorded for 5 min to establish peak current.

## Data Availability

Atomic coordinates and EM maps have been deposited in the PDB (http://www.rcsb.org) and EMDB (https://www.ebi.ac.uk/pdbe/emdb), respectively. For NaChBac in GDN, in nanodiscs, and the chimera alone and in complex with HWTX-VI, the PDB codes are 6VX3, 6VWX, 6VXO, and 6W6O, respectively; EMDB codes are EMD-21428, EMD-21425, EMD-21446, and EMD-21560, respectively.

## ACKNOWLEDGMENTS

We thank the cryo-EM facility at Princeton Imaging and Analysis Center, which is partially supported by the Princeton Center for Complex Materials, a National Science Foundation (NSF)-MRSEC program (DMR-1420541). The work was supported by grant from NIH (5R01GM130762). N.Y. is supported by the Shirley M. Tilghman endowed professorship from Princeton University.

