## Supplementary Figures S1-S7, and Table S1 for "Employing NaChBac for cryo-EM analysis of toxin action on voltage-gated Na^+^ channels in nanodisc"

### Supplementary Figures and Legends

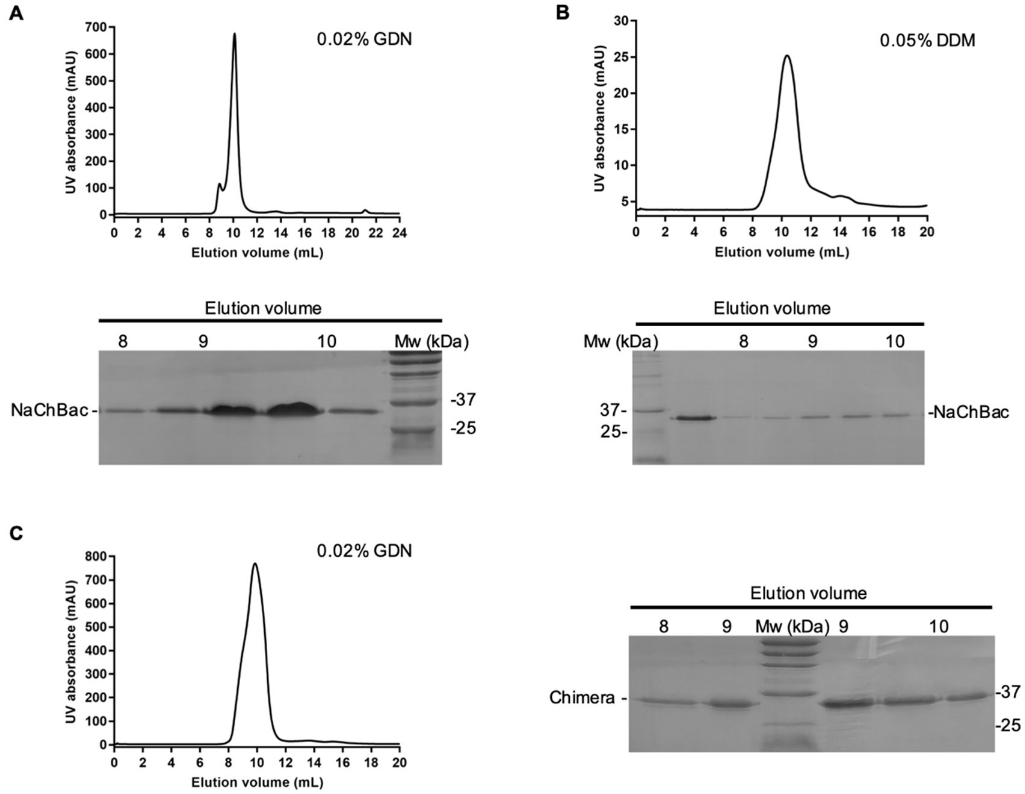


**Figure S1 | Size-exclusion chromatography (SEC) purification of NaChBac and the chimera.** (**A, B**) Representative size-exclusion chromatograms (top) and Coomassie blue-stained SDS-PAGE (bottom) for NaChBac purified in the indicated detergents. (**C**) SEC purification and Coomassie blue-stained SDS-PAGE (right) for the chimera purified in 0.02% (w/v) GDN. The pH for all three conditions was 10.0 buffered by 25 mM glycine in the presence of 150 mM NaCl.

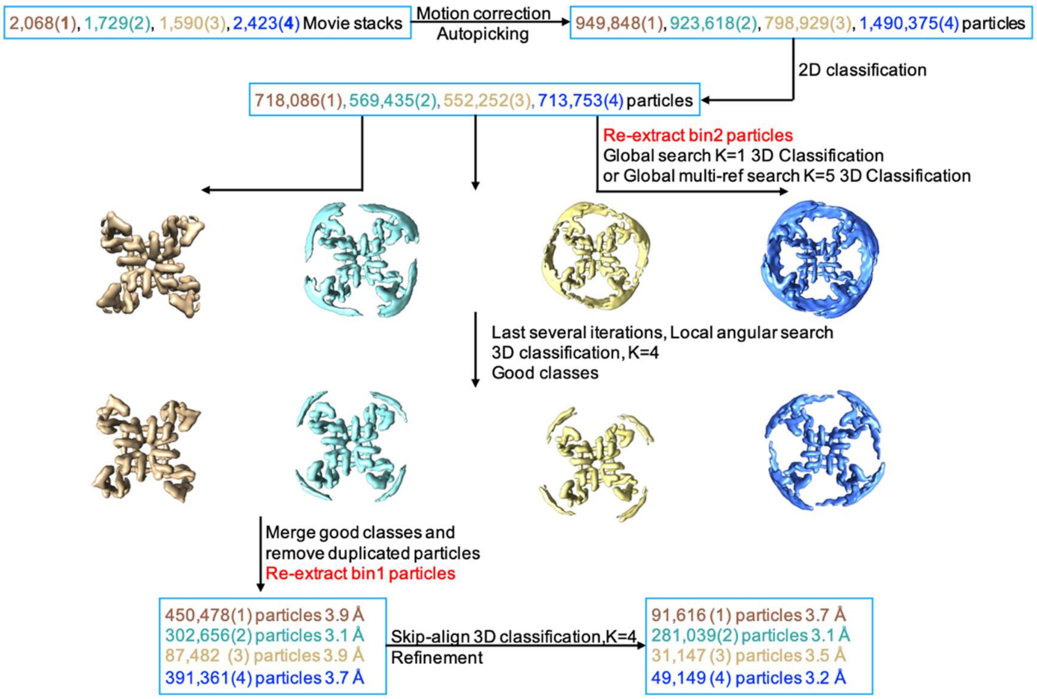


**Figure S2 | Flowchart for EM Data Processing.** Details can be found in the image processing section in Methods. From left to right or from top to bottom: (1) NaChBac in GDN, (2) NaChBac in nanodiscs, (3) chimera in nanodiscs, and (4) chimera-HWTX-IV in nanodiscs.

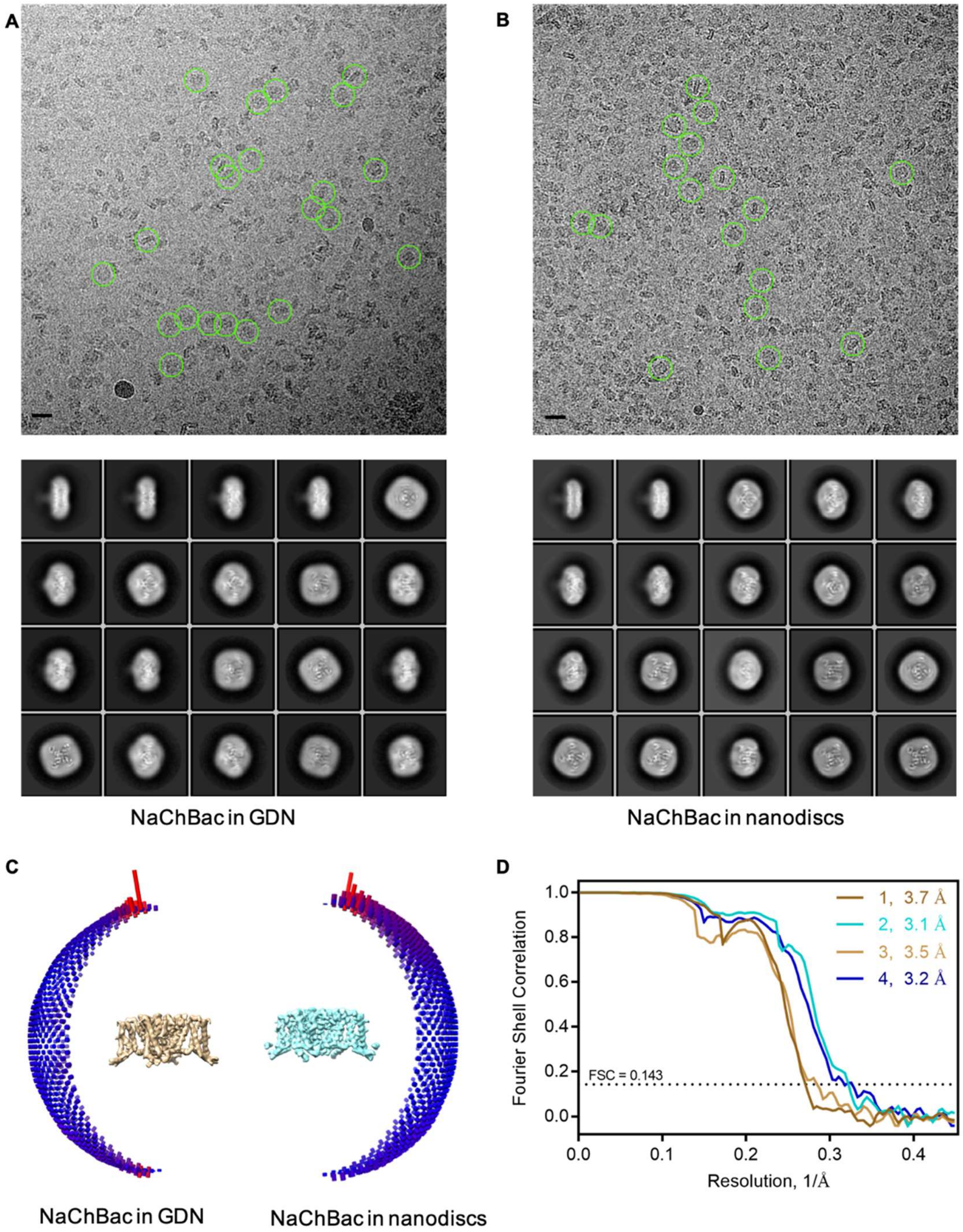


**Figure S3 | Cryo-EM data analysis.** (**A, B**) Representative micrographs and 2D class averages of NaChBac in GDN micelles (**A**) and lipid nanodiscs (**B**). Scale bar, 200 Å; box size: 260 Å; circle mask: 220 Å. (**C**) Angular distribution of the particles of the final reconstruction of NaChBac in GDN (left) and nanodiscs (right). (**D**) Gold-standard Fourier shell correlation (FSC) curves for the 3D EM reconstructions of (1) NaChBac in GDN, (2) NaChBac in nanodiscs, (3) chimera in nanodiscs, and (4) the complex between the chimera and HWTX-IV in nanodiscs.

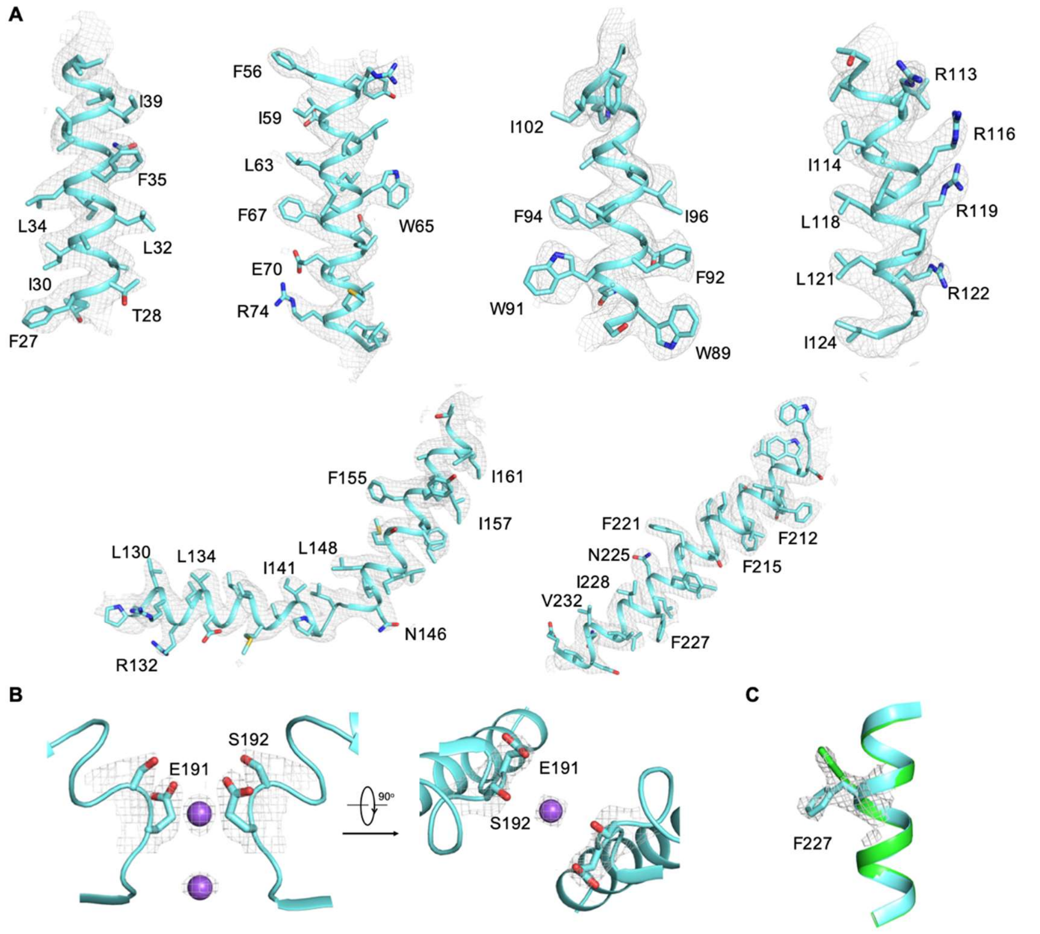


**Figure S4 | EM maps for representative segments of NaChBac in nanodiscs.** (**A**) The density of the S1-S6 segments. (**B**) The density of the selectivity filter. Two perpendicular views are shown. (**C**) The density of Phe227 shows two conformations. The "down" conformation is presented in Fig. 1C. The maps, shown as gray mesh, are contoured at 7σ and prepared in PyMol.

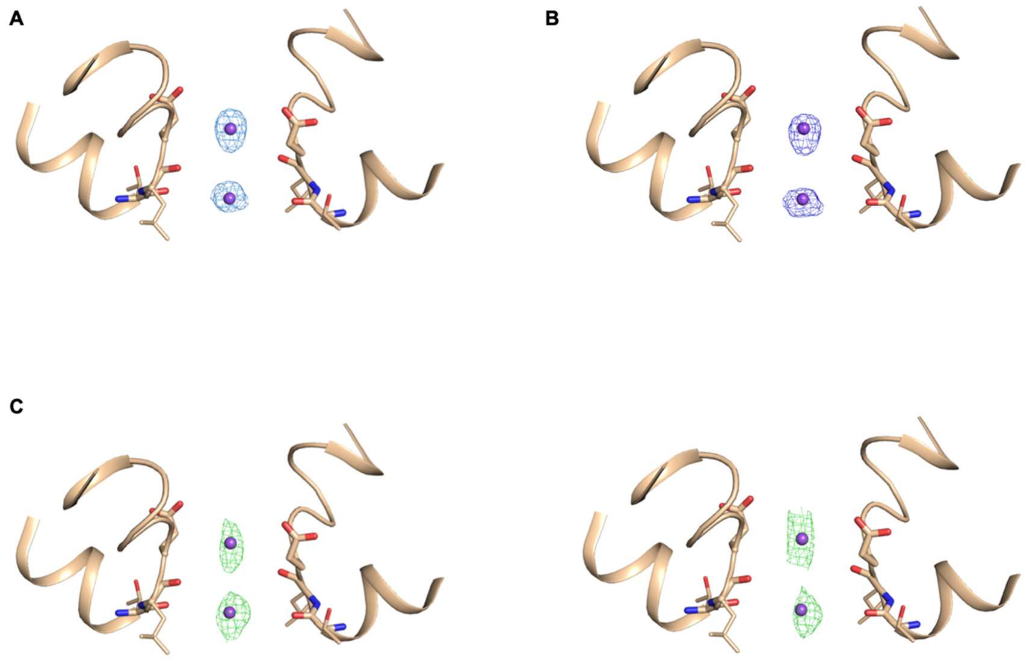


**Figure S5 | EM densities of the bound ions in the SF of NaChBac embedded in nanodiscs.** (**A, B**) The densities in the SF vestibule of nanodisc-embedded NaChBac processed with C4 (**A**) or C1 (**B**) symmetry. (**C**) Densities in the half maps. The selected particles of nanodisc-embedded NaChBac from 3D auto-refinement with C1 symmetry were randomly split into two halves, and each was subject to 3D auto-refinement. Shown here are the densities in each half map.

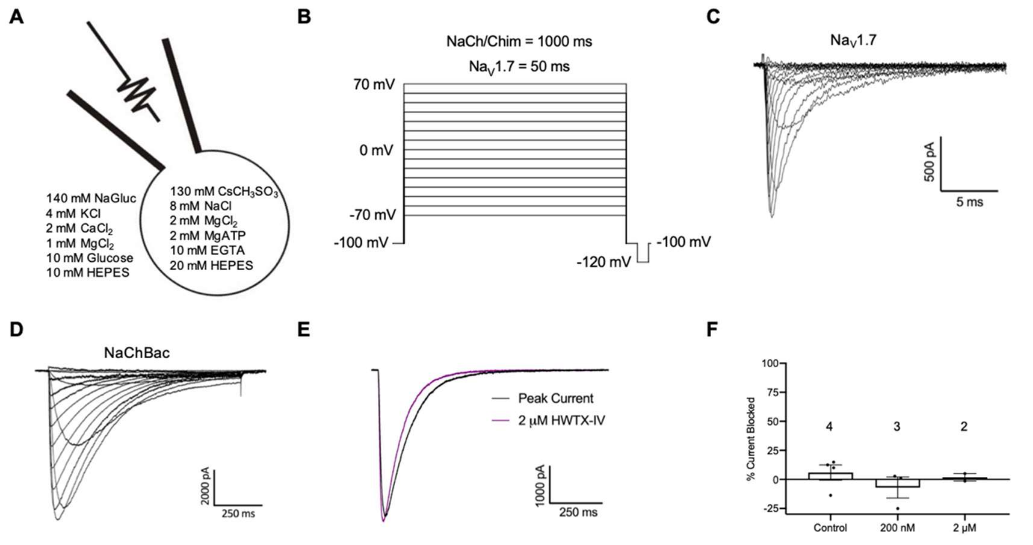


**Figure S6 | Whole-cell patch clamp recordings of exogenously expressed Na$^+$ channels**. (**A**) Bath and pipette solutions used for electrophysiological experiments. (**B**) Na$^+$ currents evoked via a step protocol from -70 mV to +70 mV in 10 mV increments. The step duration was longer for NaChBac and the chimera (1000 ms) compared to Na$_V$1.7 (50 ms). (**C, D**) Representative I-V recordings for Na$_V$1.7 (**C**) and NaChBac (**D**). (**E**) Representative single-step recording of NaChBac current before (peak current) and after 15 min perfusion with 2 μM WT HWTX-IV. Peak NaChBac current was observed within 10-15 s. (**F**) Percentage NaChBac current block (mean +/- SEM) for control (no toxin; n = 4), 200 nM WT HWTX-IV (n = 3), and 2 μM WT HWTX-IV (n = 2).

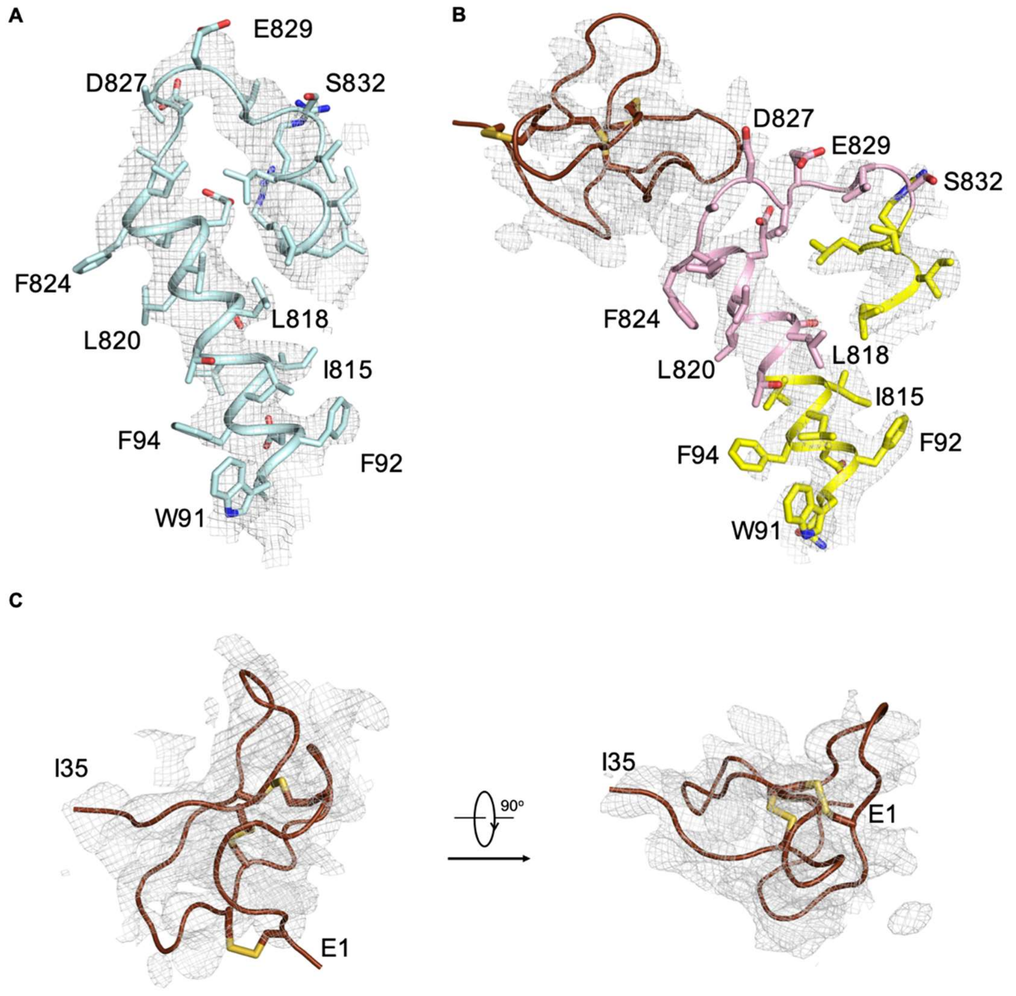


**Figure S7 | EM maps for the S3-S4 loop in the HWTX-IV-bound and the apo chimera.** (**A**) The density for the S3-S4 loop of the chimera in nanodiscs. The EM maps in panels A and B, shown as gray mesh, are both contoured at 6 σ and prepared in PyMol. (**B**) The density for the grafted S3-S4 loop of $Na_V$1.7-VSDII in the chimera and HWTX-IV. (**C**) Docking of the NMR structure (PDB code: 1MB6) of HWTX-IV into the density. The EM map, shown in two views, is contoured at 5.5 σ.

**Table S1 Summary of data collection and model statistics**

| **Dataset** | NaChBac in GDN | NaChBac | chimera | chimera-HWTX-IV |
|---|---|---|---|---|
| EM equipment | Titan Krios (Thermo Fisher Scientific Inc.) | | | |
| Voltage (kV) | 300 | | | |
| Detector | Gatan K2 Summit | | | |
| Energy filter | Gatan GIF Quantum, 20 eV slit | | | |
| Pixel size (Å) | 1.114 | | | |
| Electron dose ($e^-$ /Å$^2$) | 53 | | | |
| Defocus range (μm) | 1.5 | | | |
| # movie stacks | 2068 | 1729 | 1590 | 2423 |
| Software | Relion 3.0 | | | |
| Number of particles | 91,616 | 281,039 | 31,147 | 49,149 |
| Symmetry | C4 | | | |
| Resolution (Å) | 3.7 | 3.1 | 3.5 | 3.2 |
| Map sharpening B-factor (Å2) | -195 | -142 | -128 | -113 |
| Software | Phenix | | | |
| Cell dimensions | | | | |
| a=b=c (Å) | 267.36 | 267.36 | 267.36 | 267.36 |
| α=beta=gamma (˚) | 90 | 90 | 90 | 90 |
| Model composition | | | | |
| Protein residues | 900 | 900 | 904 | 904 |
| Ligands | 8 | 8 | 12 | 12 |
| R.m.s. deviations | | | | |
| Bonds length (Å) | 0.007 | 0.008 | 0.009 | 0.009 |
| Bonds angle (˚) | 1.01 | 1.04 | 0.77 | 0.99 |
| Ramachandran plot statistics (%) | | | | |
| Preferred | 92.60 | 91.48 | 87.39 | 83.27 |
| Allowed | 7.40 | 8.07 | 12.17 | 15.18 |
| Outlier | 0 | 0.45 | 0.45 | 1.56 |
